# Formulation of a composite nasal spray enabling enhanced surface coverage and prophylaxis of SARS-COV-2

**DOI:** 10.1101/2020.11.18.388645

**Authors:** R. J. A. Moakes, S. P. Davies, Z. Stamataki, L. M. Grover

## Abstract

Airborne pathogens pose high risks in terms of both contraction and transmission within the respiratory pathways, in particular the nasal region. Although knowledge of airborne transmission has long been known, there is little in the way of adequate intervention that can protect the individual, or even prevent further spread. This study focuses on a nasal applicant with the capacity to combat such issues, by focussing on the SARS-CoV-2 virus. Formulation of a spray containing polysaccharides known for their mucoadhesive properties was undertaken and characterised for their mechanical, spray patterns and antiviral properties. The ability to engineer key behaviours such as yielding have been shown, through systematic understanding of a composite mixture containing two polymers: gellan and λcarrageenan. Furthermore, spray systems demonstrated highly potent antiviral capacities, resulting in complete inhibition of the virus when studied for both prophylaxis and prevention of spread. Finally, a mechanism has been proposed to explain such findings. Therefore, demonstrating the first fully preventative device, targeted to protect the lining of the upper respiratory pathways.

## Introduction

Transmission of viruses occurs through 4 routes: direct contact, via physical contact with a carrier; indirect contact, interactions with contaminated objects; droplet and airborne transmission, often through coughs, sneezes and breathing; and, aerosolization, atomised virus suspended in airflow.^[1]^ Airborne transmission of respiratory pathogens, whether through droplets or atomisation, is particularly deleterious, with the virus effectively and locally delivered to the respiratory pathways. Recent work, primarily undertaken within the COVID-19 pandemic has heavily focused on providing a deeper understanding on person to person airborne transmission.^[2–5]^ During the act of coughing, turbulent air forces mucus breakup into droplets^[6]^ (*ca*. 0.62 to 15.9 μm),^[7]^ which are then expelled through the oronasal passages at flow rates of up to 12 Ls^-1^, reaching velocities of up to 30 ms^-1^.^[8]^ Unfortunately, the expelled cloud is subject to many varying parameters: speed of expulsion, droplet size and environmental effects such as air speed, resulting in effected boundaries ranging from *ca*. 0.5 to 8 m.^[8,9]^ It is likely, for this reason, that the inability to standardise transmission in this way has led to an ongoing lack of change in regard to the concepts employed to cope with such issues;^[10]^ with recommended distancing guidelines still based on the ideas portrayed by Chaplin and Wells close to a century ago.^[11,12]^ Although the epidemiology of the severe acute respiratory syndrome coronavirus 2 (SARS-CoV-2) is not yet definitive, clear indications suggest epidemiological characteristics closely linked to airborne transmittance^[13]^.

There are many airborne viruses including: influenza-, rhino-, adreno-, entero- and coronavirus. The latter, *coronaviridae* (CoVs) family, are implicated in a variety of gastrointestinal, central nervous system and respiratory diseases (MERS, SARS);^[14]^ with the latest strain, SARS-CoV-2, receiving much attention due to its devastating impact within the 2020 pandemic. SARS-CoV-2, like all coronaviruses, contains large positive-strand RNA genomes packed within a helical capsid, all housed within a phospholipid bilayer envelope formed on budding.^[15,16]^ Associated with the viral membrane are 3 main proteins: membrane and envelope proteins, associated with assembly, and spike proteins. The spike proteins, which give rise to its corona shape, are essential for virus survival, mediating entry to the host cell.^[17,18]^ Additionally, the protein also plays a crucial role in determining host range and tissue tropism, alongside being responsible for inducing many of the host immune responses.^[14]^ To date, facilitation of viral entry into a host cell is believed to arise through specific motifs within the spike protein, which strongly interact with Angiotensin-Converting Enzyme 2 (ACE2) receptors.^[19,20]^ ACE2 is known for its role in regulating oxygen/carbon dioxide transfer, commonly found within the respiratory epithelia. In particular SARS-CoV-2 has been found to target the ciliated and goblet cells,^[21]^ where subsequent viral shedding results in extensive viral loads, especially within the upper respiratory tract.^[22]^

Respired air is primarily routed through the nose. Even though the nasal passages present the highest resistance to airflow, on average *ca*. 10,000 L of air is inhaled by a healthy human per day.^[23,24]^ Only once this pathway becomes overloaded does the body switch to respiratory through the mouth.^[25][26]^ For this reason, the nasal cavity supports two major roles: climate control, creating the correct levels of humidity and air temperature; and, removal of foreign particles including dust, airborne droplets and pathogens.^[27]^ Anatomically, the nose consists of 2 cavities roughly 10 cm in length and half again in height, producing a total surface area of about 150 cm^2^.^[28]^ Inspired air flows up through the nasal vestibule (nostril) and passes through the slit-like meatus structures (inferior, middle and superior) and back through the nasopharynx. At a cellular level the majority of the cavity consists of a typical airway epithelium, comprising of 4 main cell types: basal, ciliated/non-ciliated columnar and goblet cells. The columnar cells, whether ciliated or not, are coated by microvilli. Their role, to prevent drying, supports the cilia in performing mucociliary clearance of mucins produced in the goblet cells.^[29,30]^ Additionally, the presence of cilia and microvilli drastically increases the effective surface area (*ca*. 9.6 m^2^), providing a highly efficient platform for filtration.^[31]^ Unfortunately, such large surface areas also provides greater exposure in terms of viral entry.

The airborne risk imposed not only through ventilation systems and crowds, but re-suspension of the virus from inanimate objects, including personal protective equipment,^[32]^ vociferates the need for new and novel devices that not only prevent contraction, but stop spread thereafter. This study looks to address such issues, by specifically engineering spray formulations which target the nasal passages. The emphasis on speed within such unprecedented times in terms of translating the fundamental science from lab to clinic, drives key considerations such as simplicity and proven biocompatibility. As such, colloidal composites of FDA approved polymers were studied for their application as nasal sprays. Systems were deconstructed back to their single constituents, and characterised for their mechanical, spray and antiviral properties. As such a set of design principles was determined in order to present a potential nasal spray to combat airborne pathogens, in particular SARS-CoV-2.

## Results

### Physico-mechanical behaviours of the nasal spray formulation

On application, nasal sprays directly contact the nasal mucosa lining the epithelium (Fig. 1a). Longevity of the applicant can be improved via careful choice of the polymer, promoting interaction with the mucus; known as mucoadhesion. A range of polymers known for their mucoadhesive properties (gellan, carrageenan, alginate, pectin, dextran) were screened through spray application to a 45° acetate surface (Fig. 1b). Polymers were classed for their ability to create an even coverage whilst being retained at the sprayed site. Fig. 1bii and 1biii shows typical images for several of the polymers tested, demonstrating a “good” and “poor” candidate; gellan and alginate respectively. Screening in this manner provided a means to narrow the systems down to both gellan and carrageenan going forward, with others either creating heterogeneous distributions or flowing under their own mass.

**Figure 1:**
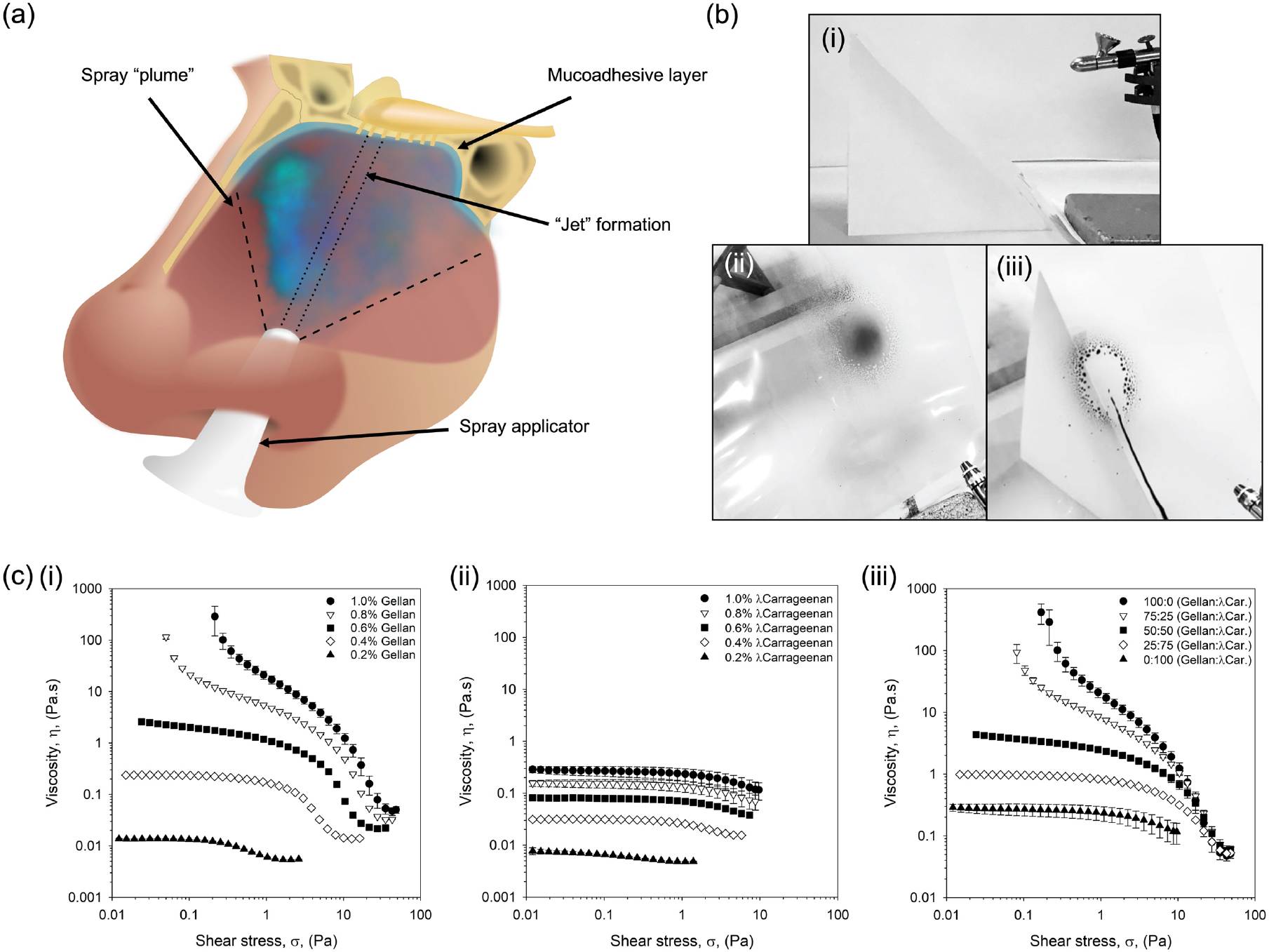
Defined nasal spray behaviours. (a) Schematic diagram demonstrating the application of a nasal spray to the nasal cavity. (b) Typical images obtained during screening of numerous mucoadhesive polymers for their ability to evenly spray and be retained on a 45° incline: (i) spray set up, (ii) gellan gum 1% (w/v) with black dye, and (iii) alginate 1% (w/v) with black dye. (c) Dynamic viscosity profiles from high to low shear stress for: (i) gellan samples with concentrations ranging from 0.2 to 1.0% (w/v), (ii) λcarrageenan samples with concentrations ranging from 0.2 to 1.0% (w/v), and (iii) composite systems of gellan: λcarrageenan at a total polymer concentration of 1% (w/v).

Flow behaviours were characterised via dynamic viscosity (from high to low shear stress), representative of the material once sprayed. Resultant profiles for the gellan were modelled demonstrating a transition from power law to Cross model, suggesting the loss of a dynamic yield stress to zero-shear viscosity as a function of the polymer concentration (Fig. 1ci). No transition was observed for the λcarrageenan systems, characterised solely by the Cross model at all polymer concentrations studied (Fig. 1cii). Zero-shear viscosity was dependent on polymer content, providing viscosities within the range of 0.27 to 0.01 mPa.s for 1.0 to 0.2% (w/v), respectively.

Viscosity curves for the composite mixtures containing both the gellan and the λcarrageenan (ratios of 100:0, 75:25, 50:50, 25:75 and 0:100) have been shown in Fig. 1ciii and Table 1. Flow behaviours for the 1% (w/v) systems showed a clear transition from material characteristics indicative of the gellan (viscosity asymptoting at low stresses), to those of the λcarrageenan (plateaued viscosities at low stresses), as the ratio of the two polymers shifted from one extreme to the other (gellan to λcarrageenan). Loss of overall viscosity was also observed as the systems shifted from high to low gellan ratios, confirmed by the reduction in consistency coefficient (K) from 3.54 to 0.03. This correlated well with the increase in rate index (n), where more gellan resulted in higher degrees of shear thinning: 0.40 to 0.82 for 100% gellan and 100% λcarrageenan, respectively. A reduction in the total polymer content to 0.4% (w/v) resulted in all mixtures characterised by the Cross model, consistent with data provided for the isolated polymers. Further reduction in the polymer concentration, to 0.2% (w/v), resulted in profiles independent on the ratio of gellan to λcarrageenan, with samples indistinguishable from each other (within error).

**Table 1:** Comparison of viscometry data. Tabulated viscometry data compiled for composite systems modelled either using the power law model (no zero-shear data provided) or Cross model (zero-shear data).

| Total Polymer<br>(% (w/v)) | Polymer Ratio<br>(Gellan:λCar.) | Zero-Shear Viscosity<br>(Pa.s) | Consistency<br>Coefficient (K) | Rate Index<br>(n) |
| --- | --- | --- | --- | --- |
| **1.0** | 100:0 | N/a | 3.544 ± 0.319 | 0.403 ± 0.004 |
| | 75:25 | N/a | 2.693 ± 0.075 | 0.491 ± 0.002 |
| | 50:50 | 4.080 ± 0.324 | 1.163 ± 0.034 | 0.543 ± 0.005 |
| | 25:75 | 0.988 ± 0.013 | 0.094 ± 0.002 | 0.727 ± 0.012 |
| | 0:100 | 0.274 ± 0.046 | 0.030 ± 0.009 | 0.821 ± 0.151 |
| **0.4** | 100:0 | 0.245 ± 0.002 | 0.065 ± 0.004 | 0.831 ± 0.003 |
| | 75:25 | 0.172 ± 0.005 | 0.060 ± 0.001 | 0.692 ± 0.054 |
| | 50:50 | 0.083 ± 0.000 | 0.020 ± 0.002 | 1.021 ± 0.084 |
| | 25:75 | 0.052 ± 0.002 | 0.016 ± 0.000 | 1.134 ± 0.029 |
| | 0:100 | 0.032 ± 0.001 | 0.013 ± 0.001 | 1.293 ± 0.051 |
| **0.2** | 100:0 | 0.014 ± 0.000 | 0.021 ± 0.000 | 1.315 ± 0.161 |
| | 75:25 | 0.009 ± 0.000 | 0.022 ± 0.000 | 1.413 ± 0.025 |
| | 50:50 | 0.010 ± 0.000 | 0.017 ± 0.003 | 1.341 ± 0.287 |
| | 25:75 | 0.007 ± 0.000 | 0.016 ± 0.007 | 1.283 ± 0.403 |
| | 0:100 | 0.007 ± 0.000 | 0.026 ± 0.001 | 1.288 ± 0.056 |

Viscometry data was used to better understand the potential residence of the spray within the nasal cavity. As such, Eq. 1 was used to predict the stress exerted on the material under gravity residing on an incline.

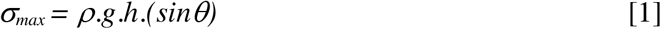

Where *ρ* is the density of the nasal spray (kg.m^-3^), *g* is the force due to gravity (9.807 m.s^-2^), *h* is the thickness of the sprayed layer (m) and θ is the inclined angle. Applying values for the polymer suspensions based on a maximum 500 μm thick sprayed layer at 45° (Eq. 2) resulted in a theoretical stress of 7 mPa.

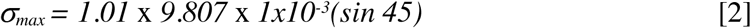

A simple force balance revealed insufficient stress under gravity to induce flow in any of the systems containing a dynamic yield stress. Indeed, even in systems described by the Cross model, the external stress due to gravity was not sufficient to move the system from its zero-shear plateau into the thinning region.

### Understanding formulation spray-ability

Application of the polymeric materials through a typical hand spray aperture has been shown in Fig. 2. Spray distributions for the single polymer systems have been demonstrated in Fig. 2a. Gellan demonstrated an inherent ability to spray forming a typical “plume” across all concentrations studied. In contrast, even at the lowest concentration, λcarrageenan systems demonstrated a degree of “jetting”, becoming more visible as the polymer concentration increased. Aspirate formation on leaving the nozzle was reflected in the distributions formed on contact with the substrate (Fig. 2b). Here, following increasing polymer, distributions became narrower with fewer satellite droplets forming around the central accumulation. A general negative correlation between %coverage and total polymer concentration was drawn, loosely fitting a linear trend (R^2^ = 0.72 and 0.62 for both gellan and λcarrageenan, respectively) (Fig. 2ci). Furthermore, it was observed that all gellan concentrations resulted in higher coverage than the λcarrageenan, demonstrating maximum and minimum %coverage of 28.5-20.7% when compared to 15.9-6.1% for the λcarrageenan systems (0.2 and 1.0% (w/v) polymer, respectively).

**Figure 2:**
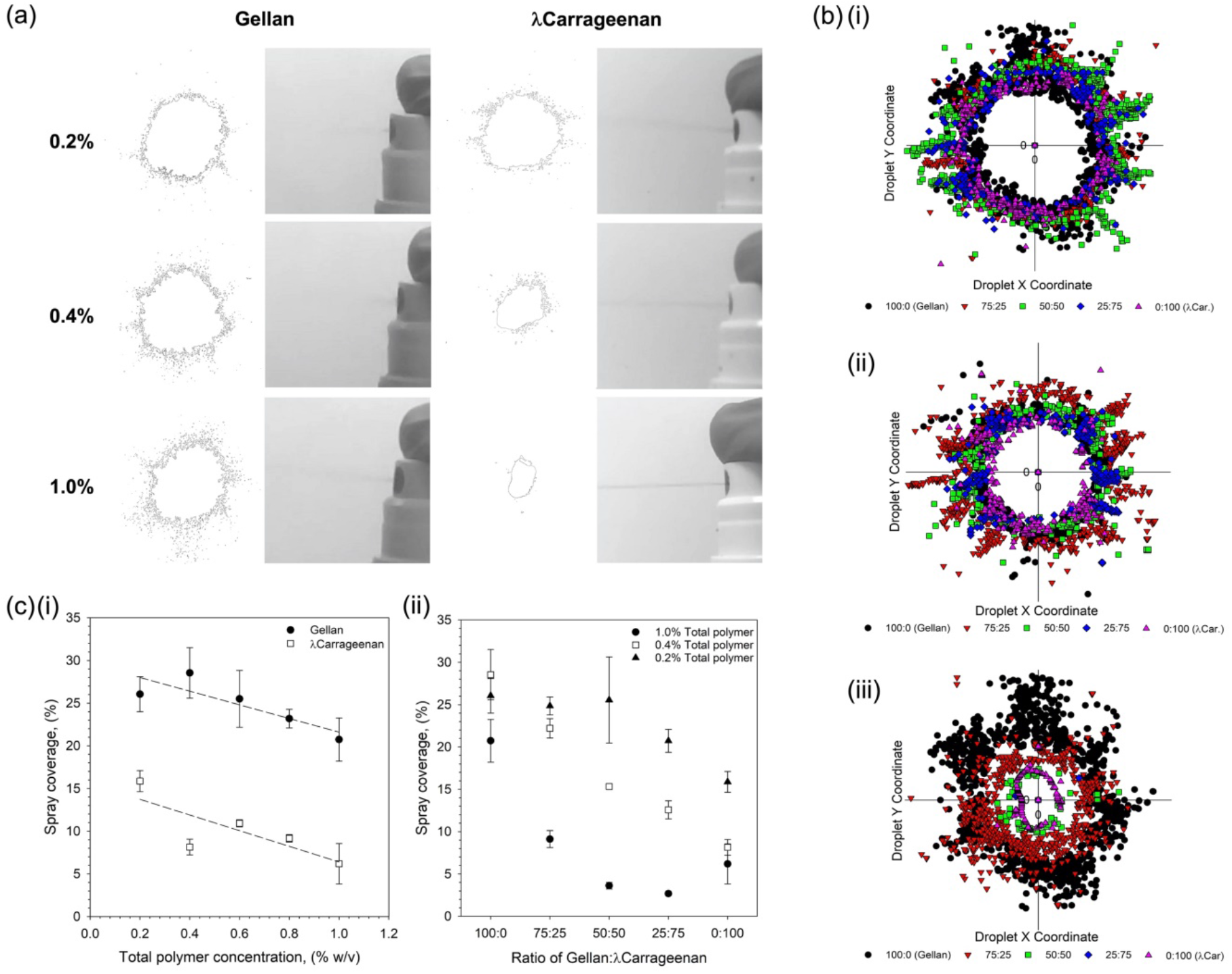
Spray-ability of polymer suspensions. (a) typical images of the spray formation as the polymer suspensions are aspirated form the applicator, alongside resulting distribution outlines for a range of polymer concentrations. (b) overlay of droplet distributions from a central point showing the reduction in spray as a function of the ratio of gellan to λcarrageenan for: (i) 0.2% (w/v) total polymer, (ii) 0.4% (w/v) total polymer, and (iii) 1% (w/v) total polymer. (c) spray coverage as determined using an imaging software for: (i) single polymer suspensions (trend lines are denoted by the dashed line with R2 values of 0.72 and 0.62 for the gellan and λcarrageenan, respectively), and (ii) composite mixtures of the gellan and λcarrageenan at either 0.2, 0.4 or 1.0% (w/v) total polymer.

The role that total and ratio of polymer play within the spray-ability of the composite systems can be clearly seen in Fig. 2b. In all instances, irrespective of total polymer concentration, a shift to smaller distributions was observed as the ratio of gellan to λcarrageen decreased. Such changes became more pronounced with total polymer, where the magnitude of change between 100% gellan to 100% λcarrageenan, followed 1.0%> 0.4%> 0.2% (w/v). such observations were mirrored in the total coverage data (Fig. 2cii). Replacing 25% of the total λcarrageenan with gellan resulted in a 4.9% and 4.4% increase in coverage, for the 0.2% and 0.4% (w/v) systems; with an initial loss in spray coverage (−3.5%) for the 1% total polymer content. Coverage was further increased to 9.0%, 14.1% and 2.9% for the 0.2, 0.4 and 1.0% (w/v) systems respectively at a ratio of 75:25 (gellan:λcarrageenan).

### In vitro analysis of the nasal spray formulations

First hit and transmission of the virus was studied *in vitro* using SARS-CoV-2 infection of Vero cells. An initial study was undertaken to determine cell compliance with the sprays over a 48 hrs incubation period (Fig. 3ai). Cell tolerance was dependent on the polymer concentration, demonstrating a 2-fold reduction in the number of living cells for both the gellan and the λcarrageenan at a dilution of 1/2. Dose-response of cell viability was linear (R^2^ = 0.96 and 0.97 for gellan and λcarrageenan, respectively), with reduced cell death as the systems became increasing more dilute.

**Figure 3:**
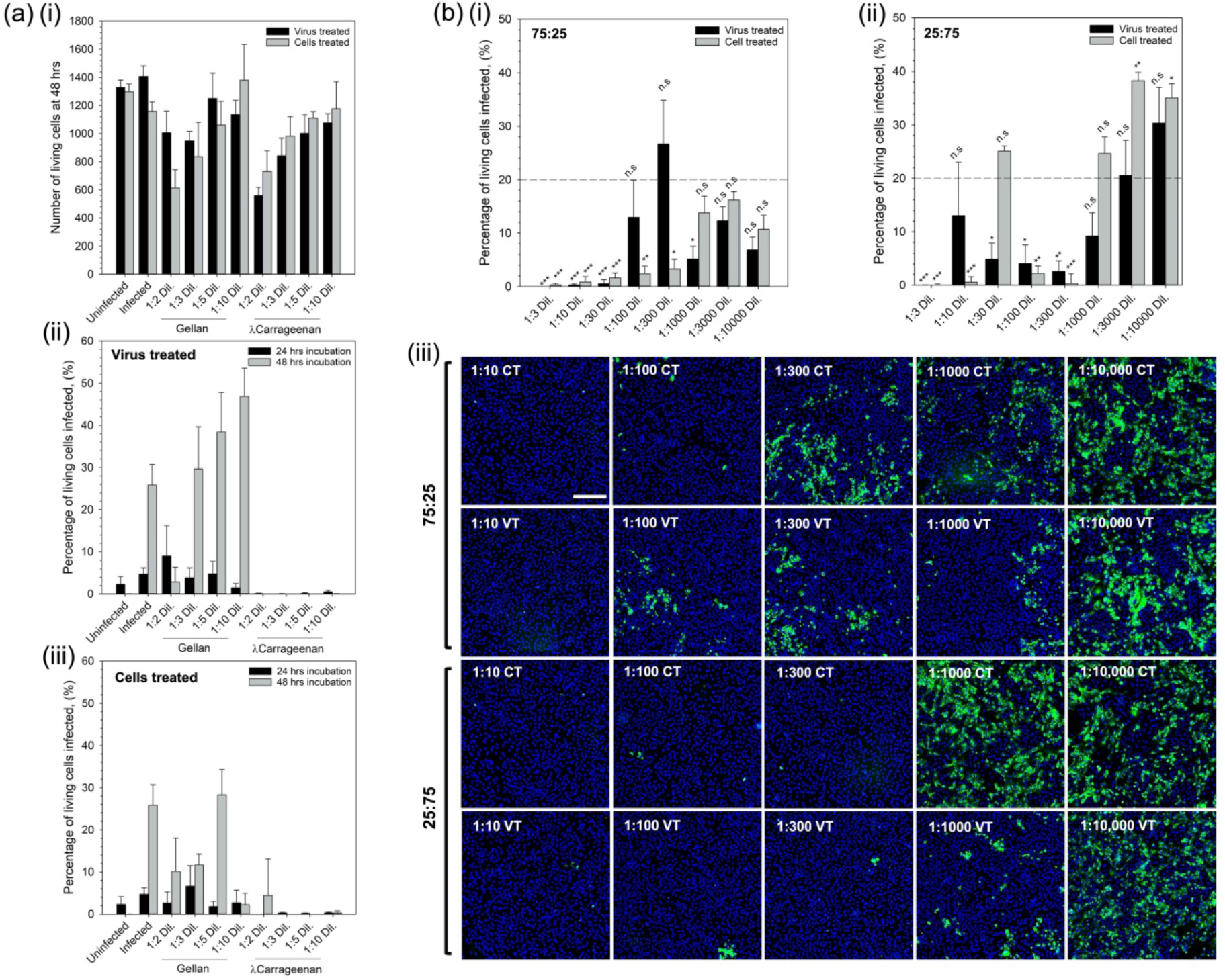
First hit and transmission analysis prevention. In vitro SARS-CoV-2 assay using vero cells to determine (a)(i) cell tolerance to the nasal sprays (live/dead analysis), (ii) degree of infection at 24 and 48 hrs for cells inoculated with the virus having undergone a pre-treatment with either the gellan or Icarrageenan spray, and (iii) degree of infection at 24 and 48 hrs for spray-treated cells inoculated with the virus. (b) degree of infection for the composite mixtures (1% (w/v) total polymer) after 48 hrs incubation having undergone either the virus treated or cell treated regimens for, (i) 75:25% gellan to λcarrageenan, or (ii) 25:75% gellan to Icarrageenan, systems, and (iii) typical fluorescent micrographs of treated systems using Hoechst staining, scale bar shows 200 μm (blue denotes non-infected and green infected cells). (n.s – not statistically different, * - p<0.05, ** - p<0.01, and *** - p<0.001)

Prevention of both contraction and/or transmission of the virus was assessed by two treatment regimens: treating the virus with the compound prior to infecting the cells (referred to as virus treated); or, by first treating the cells before introduction of the virus (referred to as cells treated). Fig. 3aii and aiii show the effect of the single polymer systems on resultant infection when treated with the virus- and cell-first regimens, respectively. It was observed that in the case of the gellan only, all dilutions resulted in infection irrespective of treatment regime after 24 hrs. Indeed after 48 hrs, such observations were exacerbated with dilutions greater than 1/3 resulting in levels of infection above the control. Interestingly, the λcarrageenan treated systems showed no signs of infection above the uninfected control at either time point, 24 or 48 hrs, irrespective of the treatment regimen.

Composite systems containing 1% total polymer at either a ratio of 75:25 or 25:75 (gellan to λcarrageenan) were also studied using the same treatment regimens over 48 hrs; data presented in Fig. 3b. Composites of a ratio 75:25 showed significant suppression of the infection (minimum of p<0.05) up to a dilution of 1/000 on comparison with the untreated control group (Fig. 3bi). In contrast, composites at a ratio of 25:75 comprising a higher proportion of λcarrageenan, demonstrated fluctuations in suppression with dilutions of 1/30, 1/1000, 1/3000 and 1/10000 all resulting in infection levels equal to or greater than the untreated control (Fig. 3bii). Comparison of the treatment regimens highlighted key differences in the ability to supress infection. Again, for the 25:75 composite it can be seen that at lower dilution factors, between 1/3 and 1/300 (with the exception of 1/30), resulted in lower average responses for cells treated prior to infection when compared to treating the virus first. However, at larger dilution factors (>1/300) it became apparent that treatment of the virus first becomes more effective. This can be seen more clearly in the images showing Hoechst stained cells, where the extent of infected cells (green – spike 2 protein staining) was much less for the virus treated groups when compared to the cell treated groups.

### Spray mechanism of inhibition

Polymer chemistry was studied in relation to the polymer type and degree of sulphation in order to better understand the mechanism of infection inhibition. Initial experiments were conducted to ascertain adherence of the polymer to the cell membrane. Staining (Alcian blue) of the sugar chains was conducted post treatment and washing. Fig. 4b shows staining intensity as a function of: polymer type, gellan and carrageenan; and, degree of sulphation along the carrageenan backbone, i and λ. Intensity data highlighted a significant difference (p<0.001) between cells treated with a 1/3 dilution of both carrageenans when compared to the cells only group. Moreover, when compared to the stained cells only group, significance remained (p<0.01). Inter-carrageenan analysis demonstrated icarrageenan to had a higher average intensity in comparison to the λcarrageenan (56.2% and 44.4%, respectively). To determine whether the degree of sulphation across the polymer backbone was important in suppression of the infection, κ−, ι− and λ–carrageenan were studied using the SARS-CoV-2 assay (Fig. 4c). It was observed that in all cases, where the cells were treated prior to being exposed to the virus, infection was lowered to below the untreated control group (p<0.001). This could not be said for the pre-treated virus, where larger dilution factors (1/1000 and 1/3000) did not statistically affect the degree of infection for both the ι- and λ-carrageenans. Additionally, no correlation could be drawn to the extent of sulphation and its ability to supress infection.

**Figure 4:**
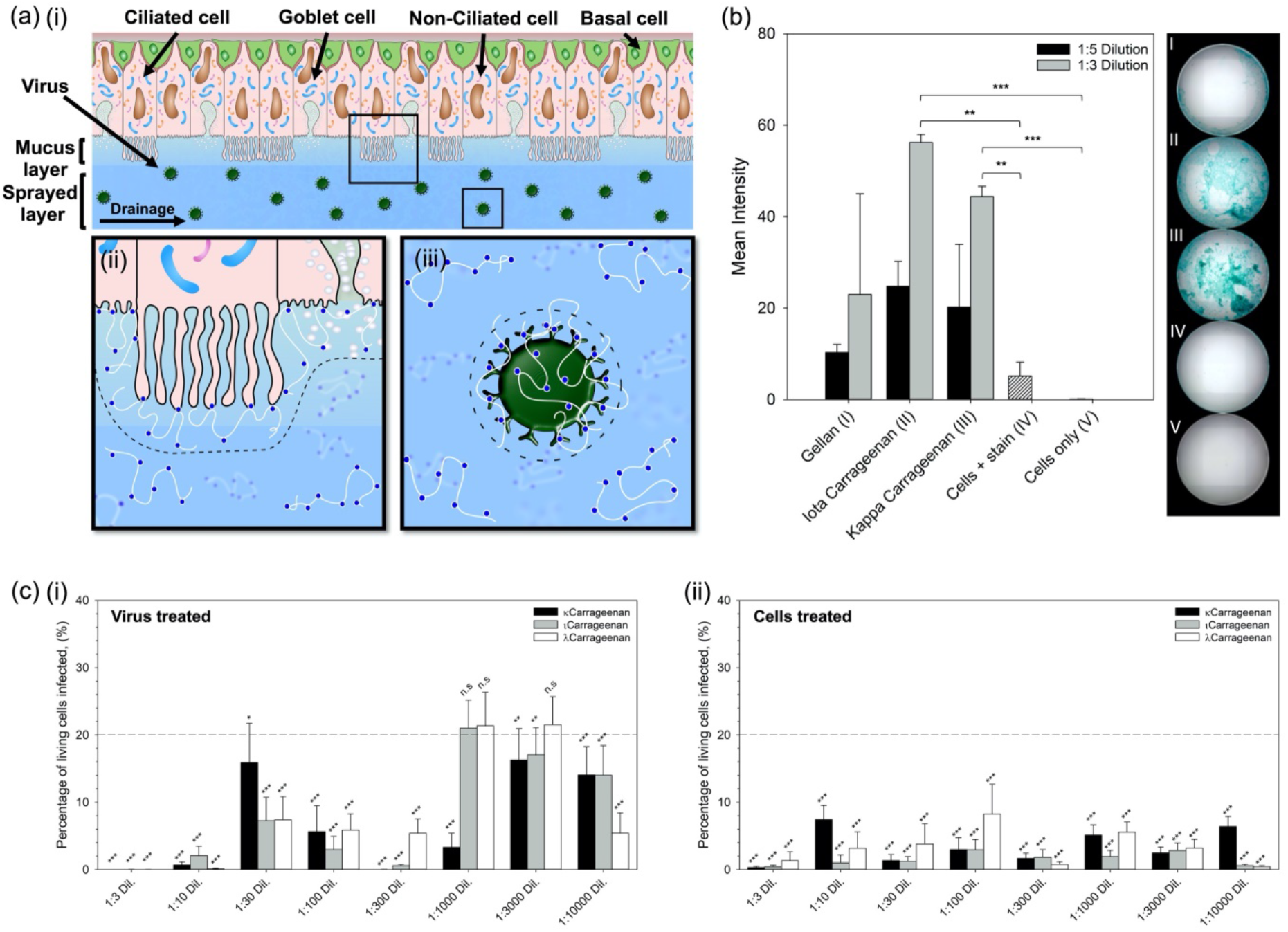
Mechanism for the inhibition of SARS-CoV-2. (a) schematic diagram showing the nasal epithelium covered in the nasal spray: (i) demonstrates potential removal of the virus via trapping within the sprayed layer and elimination through native pathways, (ii) demonstrates potential blockage of virus uptake into the cells as the polymer creates a steric barrier across the cell interface, and (iii) demonstrates potential inhibition of virus uptake by creating a steric barrier around the interface of the virus. (b) Alcian blue stain intensity for cells treated and subsequently washed with either gellan, icarrageenan or λcarrageenan. (c) In vitro SARS-CoV-2 assay using vero cells to determine levels of infection after 48 hrs for systems treated with increasingly sulphated carrageenans (κ<ι<λ), by either: (i) pre-treating the virus, or (ii) pretreating the cells. (n.s – not statistically different, * - p<0.05, ** - p<0.01, and *** - p<0.001)

## Discussion

The role that the nasal passage plays in frontline defence, filtering harmful bacteria and viruses, naturally elevates the sinonasal pathways to high risk, in terms of infection^[33]^. The need to formulate medicines/devices which can help regulate and protect this area are thus clear, however, like many regions of the body the nasal cavity poses many challenges, due to: ease of access, dynamics (native clearing mechanism) and topology (inclined surfaces or ceilings). As such, formulation engineering plays a decisive role in the design of novel therapeutics.^[34]^ The link between microstructure and material properties has long been known, ultimately driving macroscopic responses key to both function (delivery/retention/ADME – absorption, distribution, metabolism and elimination) and the end user (ability to administer/patient compliance). Through a microstructural design approach, the interplay between areas such as raw materials and processing can be manipulated to engineer defined characteristics. In the case of a nasal spray elements such as mucoadhesion, longevity, coverage and controlled delivery/prophylaxis need to be considered. Polysaccharides provide perfect polymeric candidates within biological applications, as the natural polymers often demonstrate biocompatibility and hold FDA approval; significantly reducing risk, time and costs throughout the translational process.

A simple screening process to determine the ability to evenly spray across an inclined substrate narrowed potential candidates down to both gellan gum (low acyl) and carrageenan (λ). In addition to biocompatibility, their long chains (often 100s kDa) and charged side groups (_-_COO^-^, _-_SO_3_^-^) provide inherently strong mucoadhesion through polymer entanglement, ionic interactions and weaker van der Waals interactions with the mucus layer.^[35,36]^ Such interactions with the mucosa provide high retention to the surface and a mechanism of clearance, becoming transported by the cilia out of the paranasal sinuses to the pharynx and eventually into the oesophagus.^[29,30]^

Enhancing longevity within the nasal cavity can also be sought enhanced viscosity and resultant reduction in flow/clearance. The role that gellan and carrageenan play within viscosity modification and related sensory attributes has been well established within the food industry.^[37]^ Again, owing to their long polymeric chains and chemistries along their backbone both gellan and λcarrageenan are able to structure large volumes of water. This, accompanying polymer-polymer entanglements, ultimately drives increases in viscosity,^[38,39]^ with higher polymer concentrations resulting in more viscous suspensions: until sufficiently concentrated in the case of the gellan (>0.8% (w/v)), providing the evolution of a dynamic yield stress. Again, yielding behaviour can be used to enhance application and slow clearance, as the gravitational stress is insufficient to cause rupture of the film formed post-spraying; where the film height can be estimated as a function of a typical nasal dosage (25 – 200 μl)^[40]^ over a surface area *ca*. 5 cm^2^.^[31]^

The large surface areas in the nasal cavity provides the ability to process large volumes of air (up to 35 Lmin^-1^ before switching to oronasal breathing), within a total volume of *ca*. 15 ml.^[31,40]^ However, the large nasal area presents a challenge to uniformly coat. Coverage of the polymer systems demonstrated clear correlations between both the type of polymer and the concentration of polymer used. Gellan systems demonstrated high levels of coverage across all concentrations studied, suggesting an ideal candidate for nasal spray application. Interestingly, λcarrageenan even though characterised by a lower viscosity, resulted in poor overall coverage whilst still maintaining concentration dependency. Such changes were a direct result of a shift from plume to jet formation, with gellan resulting in much faster rates of jet destabilisation in comparison to the λcarrageenan. Spray behaviours comply with literature, suggesting that large surface tensions as opposed to viscosity are required to force droplet breakup, relative to the density of the surrounding medium.^[41]^ As a result, the persistence of a jet negatively effects patient compliance, not only providing poor coverage, but eliciting unwarranted irritation on contact with the nasal wall.^[42,43]^

To maintain the advantage of λcarrageenan’s intrinsic anti-viral capacity^[44–46]^, formulation of a composite mixture containing increasing amounts of gellan to λcarrageenan allowed for optimisation of the nasal therapy. Careful control over the two polymers provided a means to engineer enhanced λcarrageenan spray-ability. Interchanging 25% of the initial λcarrageenan with gellan saw an increasing in the total area coated up to *ca*. 35% of its initial coverage. This was further increased to *ca*. 63% on replacement of 75% of the initial polymer. In addition to tailorable spray profiles, composite systems demonstrated a means to formulate sprays containing λcarrageenan with both yielding and augmented viscosities, not possible with the λcarrageenan alone. Data showed that the formation of intermediate products, from 100% λcarrageenan to 100% gellan, transitioned in behaviour governed primarily by the dominating polymer. As such, it was possible to detail a set of design principles that can be used to formulate various sprays, with desired mechanical properties.

Cytotoxicity and anti-viral activity of the nasal treatments were assessed using a relevant enveloped virus, SARS-CoV-2, and their current gold-standard model for infection (Vero cells). Initial cytotoxicity studies revealed a degree of cell death when cultured in the presence of both the gellan and λcarrageenan. Such reductions in cell numbers here are thought to be a consequence of osmotic stress, with high concentrations of the sugars resulting in cell shock.^[47]^ The plethora of literature demonstrating the combability and use of such polysaccharides in pharma and biomaterials^[48]^ might suggest that such observations are indeed an artifact of two-dimensional cell culture, as opposed to inherent toxicity.

First hit and ability to prevent viral transmission was assessed using two main treatment regimens. Firstly, prophylaxis was assessed through application of the spray onto the cells prior to infection. Gellan systems showed limited ability to suppression the SARS-CoV-2 virus, whereas, λcarrageenan demonstrated complete inhibition over 48 hrs. Pre-treatment of the virus, representing the ability to nullify the virus preventing transmission mirrored prophylaxis data, with complete inhibition; supporting previous acknowledgements that λcarrageenan provides enhanced anti-viral capacities.^[49]^ Composites again provided the ability to accommodate synergistic behaviours from both gellan and λcarrageenan: enhanced mechanical responses towards spraying and anti-viral activity. Interestingly, systems containing a greater proportion of gellan outperformed the λcarrageenan dominated system; an unexpected outcome based on the single polymer data. Indeed, the spray was highly potent with dose-dependency demonstrating significant prevention/reduction of infection up to 30- and 300-fold dilutions for the virus and cell treatments, respectively.

It is suggested that inhibition of the infection results through 3 main mechanisms: formation of a steric barrier at the cell interface, adsorption of the polymer to the virus, and/or physical entrapment of the virus in the sprayed layer. It is proposed that polymer adsorption is facilitated through charge-charge interactions at the cell and virus membrane. Although both anionic in nature, the contrast in virus inhibition infers that the carrageenan’s sulphate chemistry drives anchoring of the polymer to the substrate surface; likely through the formation of di-sulphide bridges with cationic membrane polysaccharides and/or proteins. The polymer thus provides a physical role, expanding the hydrodynamic volume around the cell/virus and preventing close proximity.^[50]^ Even though the role that the negatively charged sulphate groups play in the ability to adsorb to the bio-interface, it is unclear from the data whether a link between the degree of sulphation and suppression of infection exists. Although not significant in the role of coating, gellan does demonstrate its applicability when considering prophylaxis through entrapment and elimination. The ability to engineer high viscosities and yielding behaviour at this point becomes key, proportionally slowing diffusivity, as described by the Stokes-Einstein relation.^[51]^ To this end, diffusion of the virus towards the host cells can be hampered within timescales associated with typical nasal clearance.^[52]^ In reality a combination of the 3 proposed mechanisms is likely to occur. To this end, physical entrapment is suggested to provide a first means of defence, simultaneously resulting in a secondary defence where cells and virus become coated. Thus, any virus particles having migrated to the cell interface are already inhibited to uptake. Likewise, the formation of new viruses as a result of shedding, become incapacitated. This combinatorial approach, coupled with the highly potent anti-viral capacity of the carrageenan towards SARS-CoV-2, provides a powerful spray device with the capacity to prevent both contraction and transmission.

## Conclusions

As the primary mode of transmission for airborne viruses is uptake through the respiratory tract, the nasal passage poses one of the largest risk factors to contraction. Although it is well known that the nose filters 1000s of litres of air daily, there is little in the way of preventative measures to ensure protection to infection. This study has demonstrated the formulation of a potent antiviral nasal spray, with not only prophylactic capacity, but the ability to prevent viral transmission. Its ability to completely inhibit infection is derived from the chemistry (sulphated polymer backbone) of the active polymer, λcarrageenan. Spray characteristics were engineered through the production of a composite, where a set of design rules were understood to allow for manipulation over the material behaviours: spray coverage, viscosity and yielding behaviour. Furthermore, understanding the role of each polymer in the composite allowed for a preventative mechanism, using the synergy of both material and antiviral properties to coat the biological interfaces, prevent viral uptake by host cells, and eliminate through native clearance pathways. As such, this work presents a potential device with the capacity to specifically target infection within the nasal cavity.

## Materials and methods

### Materials

Sodium alginate (medium viscosity), pectin from citrus peel, dextran (Mw *ca*. 20 kDa), kcarrageenan, icarrageenan, λcarrageenan, PBS, heat inactivated FBS, Penicillin/streptomycin, Alcian blue (8GX) were all purchased from Sigma Life Science, UK; Gellan gum (CG-LA) was purchased from CP Kelco; TrypLE Express 1x was purchased from Fisher Scientific; Black dye (Parker); Type 1 water (Milli-Q, Merck Millipore).

**Single-Component systems –** colloidal suspensions were prepared through the addition of polymer (0.2 to 1.0% (w/v)) to a dilute PBS (5% v) solution. Once added, the systems were vigorously mixed and left to fully hydrate for 24 hrs. All samples were kept at ambient temperature (*ca*. 20 °C) until further used.

**Multi-Component systems –** composite mixtures were prepared by first weighing out ratios of polymer (75:25, 50:50, 25:75 – gellan gum (low acyl) to λcarrageenan), and thoroughly mixing. Powdered mixtures (0.2, 0.4 and 1.0% (w/v) total polymer concentration) were then added to a dilute PBS (5% v) solution, vigorously mixed and left to fully hydrate for 24 hrs. All samples were kept at ambient temperature (*ca*. 20 °C) until further used.

## Screening

Polymer screening was conducted using an airbrush (750 μm aperture) coupled to an oil-free compressor (Badger, US), set to 1 bar. Test material (0.9 ml) was mixed with black dye (0.1 ml) and sprayed across an acetate sheet set to a 45° incline. The airbrush was then cleaned using a succession of 70% ethanol and water. Spray distributions were visually analysed for homogeneity and retention.

## Rheology

Viscometric analysis was undertaken on a rotational rheometer (Kinexus Ultra, Netzsch Geratebeu GmbH, DE) fitted with a cone and plate (4°, 40 mm diameter) geometry. Tests were conducted at 25 °C, under stress control. Dynamic viscosity was analysed by reduction of the shear stress from a maximum of 100 to 0.001 Pa (dependent on test material to prevent expulsion from the gap at lower viscosities) over a 2 mins ramp time. Kinexus software was used to characterise the flow profiles using both power law and Cross models.

## Spray-ability

Test material was first mixed with black dye (0.1% v) and thoroughly shaken to provide a homogenous mixture. A typical handheld applicator (Adelphi, UK) was used to vertically spray a paper recipient. Sprayed distributions were allowed to dry in air (no blotting effects observed) and scanned at 600 DPI (greyscale). Image files were processed using an image package (ImageJ), where they were initially cropped to a 2000 by 2000 px box visually centred around the spray pattern. Standard thresholding was applied to all images, and scale corrected equating 2000 px to 100%. Droplet analysis was conducted, and total coverage determined as a percentage of the whole image. Distributions were recorded as x/y co-ordinates and plotted relative to the central droplet.

## Infection/transmission

Vero cell were washed with PBS, dislodged with 0.25% Trypsin-EDTA (Sigma life sciences) and seeded into 96-well imaging plates (Greiner) at a density of 10^4^ cells per well in culture media (DMEM containing 10% FBS, 1% Penicillin and Streptomycin, 1% L-Glutamine and 1% non-essential amino acids). Cells were incubated for 24 hrs to allow time for adherence. Virus or cells were treated with polymeric solutions, diluted in media, 1 hr prior to infections. Cells were subsequently infected with SARS-CoV-2 virus England 2 stock 10^6^ IUml^-1^ (kind gift from Christine Bruce, Public health England) diluted 1/150 in culture media. Cells were fixed in ice-cold MeOH after infection. Cells were then washed in PBS and stained with rabbit anti-SARS-CoV-2 spike protein, subunit 1 (The Native Antigen Company), followed by Alexa Fluor 555-conjugated goat anti-rabbit IgG secondary antibody (Invitrogen, Thermofisher). Cell nuclei were visualised with Hoechst 33342 (Thermofisher). Cells were washed with PBS and then imaged and analysed using a ThermoScientific CellInsight CX5 High-Content Screening (HCS) platform. Infected cells were scored by perinuclear fluorescence above a set threshold determined by positive (untreated) and negative (uninfected) controls.

## Cell binding

**Preparation of cells –** Vero cells were expanded in T75 flasks, washed with PBS (5 ml) and removed using TrypLE (2.5 ml). The cells were then re-suspended in complete media and seeded in to 96 well plates (10,000 cells per well). Cells were left to attach over the subsequent 24 hrs prior to treatment.

**Cell treatment –** cells were washed (3 times) with PBS and final washing removed. Test material was diluted to either 1/3 or 1/5 and placed over the cells (200 μl) (controls were treated with equal volumes of PBS). Cells were incubated for 30 minutes prior to washing (3 times) with PBS. Cells were subsequently stained with Alcian blue (0.1%) for 30 minutes, before a final wash in PBS to remove residual stain. PBS was then added (200 μl) and wells imaged.

**Cell Imaging –** cells were imaged using a Cytation 5M automated microplate imager. Wells were images in bright field using a x4 optical lens focused on the centre of each well. Wells were divided into a 6 x 4 matrix and stitched together retrospectively. Images were then cropped to the well diameter using a software package (ImageJ) and colour thresholding standardised and analysed for mean intensity.

## Statistics

In all experiments data presented is an average of at least triplicates with error portrayed as the 95% confidence interval. Significance was determined by first assessing data for normality. Where normally distributed, paired t-tests were conducted comparing the treatment group to the untreated control. If the normality test failed, comparison was made on ranks using the Mann-Whitney. Significance has been shown on plots using the following notation: n.s – not statistically different; * - p<0.05; ** - p<0.01; and, *** - p<0.001.

## Acknowledgments

The authors would like to extend thanks to: Stephen Priestnall for useful discussions; Nicholas Barnes and Vasanthy Vigneswara of the University of Birmingham Clinical Sciences for help undertaking elements of the cell work; and, Christine Bruce of Public Health England for kindly providing the SARS-CoV-2 virus.

